# Genetic potential for denitrification in SAR11 and other ubiquitous lineages of marine bacteria isolated from the northern Benguela Upwelling System

**DOI:** 10.1101/2025.06.18.660415

**Authors:** Robert M. Morris, Timothy E. Mattes, Kunmanee Bubphamanee, Mike C. Sadler, Luke A. Calderaro, Jen Karolewski, Karen L. Casciotti, Olivia U. Mason

## Abstract

Denitrification is a microbial process that leads to nitrogen loss from marine oxygen minimum zones (OMZs). The complete metabolic pathway for denitrification reduces nitrate to dinitrogen gas in four sequential steps. Many anaerobic and facultatively anaerobic bacteria are capable of the initial step of nitrate reduction to nitrite. Far fewer reduce nitrite to nitric oxide, nitrous oxide, or dinitrogen gas. In this study, we cultured and sequenced the complete genomes of 24 bacteria isolated from low dissolved oxygen waters (DO = 24 µM) in the northern Benguela Upwelling System (nBUS) OMZ to identify facultatively anaerobic bacteria with the genetic potential to contribute to denitrification. Most of the isolates obtained from the nBUS have denitrification genes (79%). They include several new species in the order *Pelagibacterales* (SAR11), as well as representatives from a previously uncultured family of *Arenicellales* (UBA868), a previously uncultured genus of *Paracoccaceae*, and a previously undescribed genus of *Porticoccaceae*. All ten nBUS SAR11 have a previously unidentified genomic region that codes for a copper-containing nitrite reductase (*nirK*), suggesting that they have the potential to contribute to nitrogen loss by respiring nitrite to nitric oxide.

**Significance Statement:** Vast genomic diversity has confounded sequencing efforts to identity the potential for marine bacteria to contribute to denitrification in marine oxygen minimum zones (OMZs). To identify facultatively anaerobic microbes with the genetic potential to contribute to marine nitrogen loss, we cultured and sequenced the complete genomes of bacteria from low-oxygen waters of the northern Benguela Upwelling System. Complete genomes allowed for a comprehensive analysis of denitrification. Most bacterial isolates, including all ten SAR11, have the genetic potential to contribute to denitrification. This suggests that some of the most abundant lineages of marine bacteria in the oceans are adapted to anoxic conditions in OMZs and may have a more direct role in marine nitrogen loss than previously suspected.

## Introduction

In marine oxygen minimum zones (OMZs), the concentrations of dissolved oxygen (DO) can fall to less than 10 nanomoles/L (Bertagnolli and Stewart, 2018). Although they account for only 1% of the ocean by volume, microbial processes in OMZ catalyze 30-50% of marine nitrogen loss (Codispoti, 2010; Lam and Kuypers, 2011). This outsized impact on the marine nitrogen budget is due largely to the anaerobic microbial processes of denitrification and anaerobic ammonia oxidation (anammox) that occur in these low oxygen waters (Kuypers et al., 2005; Lam et al., 2009; Ward et al., 2009).

Denitrification is the stepwise reduction of nitrate (NO_3_ ^-^) to nitrite (NO ^-^), nitric oxide (NO), nitrous oxide (N_2_O), and di-nitrogen gas (N_2_) (Zumft, 1997). Each step in the denitrification pathway requires a specific enzyme, and it is common for microbes to possess some, but not all, of the required enzymes for denitrification (Pold et al., 2025; Zhang et al., 2023). While many anaerobic or facultatively anaerobic bacteria can respire nitrate, fewer have the genetic potential to reduce nitrite to a gas (NO, N_2_O, or N_2_). This incomplete denitrification can lead to the accumulation of intermediates in anoxic waters and fuel competition between microbes specializing in components of the denitrification pathway (Sun et al., 2024). Low oxygen waters with these microbes can be either a source or sink of N_2_O, Depending on environmental conditions and interactions among microbial community members.

The discovery that SAR11, one the most abundant lineage of bacteria in the oceans (Morris et al., 2002), have and express genes to reduce nitrate to nitrite greatly expanded their known range beyond the oxygenated ocean and raised their perceived importance in anoxic waters of marine OMZs (Tsementzi et al., 2016; Ruiz-Perez et al., 2021). Evidence that *Pelagibacterales* (SAR11) respire nitrate and produce nitrite in the Eastern Tropical North Pacific (ETNP) OMZ also linked their activities to the dominant nitrogen loss processes in the oceans: denitrification and anammox. However, less is known about the genetic potential and distribution of facultatively anaerobic SAR11 in other OMZs.

The Benguela Upwelling System (BUS) is an important and relatively understudied OMZ in the South Atlantic Ocean. Several studies have documented significant production of nitrous oxide within the low-oxygen waters of the BUS (Frame et al., 2014; Arévalo-Martinez et al., 2019; Sabbaghzadeh et al., 2021; Dangl et al., 2025), but the microorganisms responsible for this nitrogen loss, particularly through denitrification, are still largely unknown (Dangl et al., 2025). Sulfur oxidizing γ-proteobacteria from the family *Thioglobaceae* (SUP05) have the potential to respire nitrogen oxides (reviewed in Morris and Speitz, 2022) and often dominate marine microbial communities in the BUS (Lavik et al., 2009). Many other heterotrophic bacteria that are facultatively anaerobic can also respire nitrogen oxides. To identify the potential for bacteria to contribute to denitrification in the BUS, we cultured and sequenced the complete genomes of 24 bacteria from low dissolved oxygen waters (24 μM) in the northern BUS (nBUS) OMZ.

## Results and Discussion

### Diversity of Denitrification Genes in Bacteria from the nBUS

Marine bacteria isolated from the nBUS include diverse members of the *Alphaproteobacteria*, *Gammaproteobacteria*, and *Bacteroidia* (**Figure 1**). The isolates have circular chromosomes with genomes that range in size from 1.3 to 4.6 Mbp and have GC contents between 29.6 to 52.5% (**Supplementary Table S1, Supplementary Material**). Most (19/24) have the genetic potential to carry out at least one step in the denitrification pathway. Only one isolate, *Pseudopelagicola* nBUS_20, has a full set of genes for complete denitrification, suggesting that it has the potential to reduce NO_3_ ^-^ to N_2_ (**Figure 1, Supplementary Table S2, Supplementary Material**). The complete denitrification pathway in nBUS_20 includes both periplasmic *napAB* and respiratory *narGHI* dissimilatory nitrate reductase genes, as well as a *nirS* nitrite reductase, *norC* nitric oxide reductase, and *nosZ* nitrous oxide reductase. The genome of the only *Bacteroidia* isolate, *Tenacibaculum* nBUS_03, codes for a single denitrification gene, NorC nitric oxide reductase.

**Figure 1.**
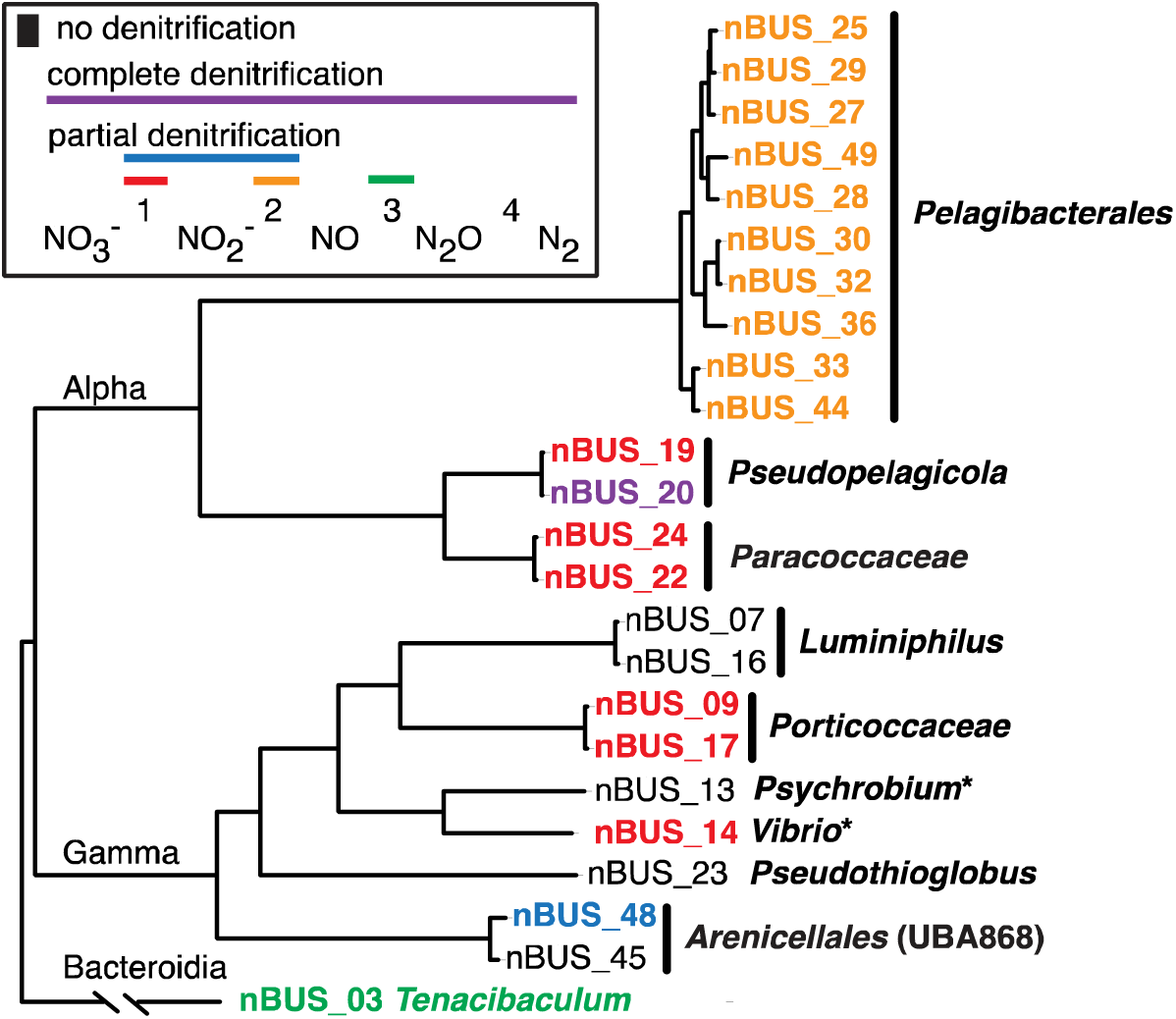
Genetic potential for denitrification in the genomes of bacteria isolated from the nBUS. Key denitrification genes include those for nitrate reductases (*napAB* and *narGHI*), nitrite reductase (*nirK*), nitric oxide reductase (*norC*), and nitrous oxide reductase (*nosZ*). Asterisks indicate the presence of assimilatory *nasC* and *nirBD* genes. Genes correspond to KEGG orthology numbers are listed under gene_id in Tables S2 and S4: *napAB* (K02567, K02568); *narGHI* (K00370, K00371, K00374); *nirKBD* (K00368, K00362, K00363); *norC* (K02305); *nosZ* (K00376); and *nasC* (K00372).

Genes to respire only nitrate and nitrite were identified in one genome and to respire only nitrate were identified in seven genomes. Isolate nBUS_48 is from a previously uncultured family in the order *Arenicellales*, UBA868 (Baltar et al., 2023), which encodes *narGHI* and *nirK* dissimilatory nitrate and nitrite reductase genes. The *Vibrio* nBUS_14 genome encodes *napAB* dissimilatory nitrate reductase genes, as well as key components of assimilatory nitrate and nitrite reduction pathways, including the cytoplasmic *nasC* and *nirBD* genes, respectively.

*Psychrobium* nBUS_13, has the cytoplasmic *nasC* and *nirBD* genes. The NirBD nitrite reductases have known roles in fermentative dissimilatory nitrate reduction to ammonium (DNRA) (Pandey et al., 2020), but they are generally linked to assimilatory nitrite reduction (Besson et al., 2022). Other key components of assimilatory nitrate reduction (*nasB*) and fermentative DNRA (*narGHI*) were not identified in these isolates, which suggests that the combination of cytoplasmic systems found in nBUS_13 and nBUS_14 may represent an alternative pathway for nitrate reduction to ammonium.

Genes for nitrate respiration were broadly distributed in the nBUS bacterial community. For example, three *Alphaproteobacteria* and three *Gammaproteobacteria* were identified with the genetic potential to respire nitrate (**Figure 1**). The *Alpharoteobacteria* include *Pseudopelagicola* nBUS_19 and two representatives from a previously uncultured genus in the family *Paracoccaceae*, nBUS_22 and nBUS_24. The *Gammaproteobacteria* include two representatives from an undescribed genus in the family *Porticoccaceae* (Stingl et al., 2007), nBUS_09 and nBUS_17, and *Vibrio* nBUS_14.

### A Conserved NirK-cbb3 Genomic Region in nBUS SAR11

Ten new SAR11 species and subspecies were isolated and sequenced from the nBUS (**Figure 2**). They are members of subclade 1a.III.IV (**Figure 2a, Supplemental Material**), which is a lineage that was recently assigned the genus *Proprepelagibacter* (Freel et al., 2024). Phylogenomic and whole genome average nucleotide identity (ANI) analyses indicate that the nBUS SAR11 represent six new species and four new subspecies, with pairwise ANI values that range from 0.85-0.98 (**Supplemental Table S3, Supplemental Material**). The most striking feature in the nBUS SAR11 genomes is that all ten have a conserved genomic region containing a copper-containing nitrite reductase (*nirK*) and components of cbb3-type cytochrome c oxidases (*cco*) that are not present in the complete genomes of SAR11 cultured from oxygenated waters in the North Pacific, North Atlantic, and Indian oceans (**Figure 2b; Supplemental Table S4, Supplementary Material**). Notably, the combined percent GC for *nirK*-cbb3-type cytochrome c oxidase genes in all ten SAR11 is 35.7%, which is higher than the average for the genomes (30.1%). This suggests that the *nirK*-*cbb3* genomic region in SAR11 is a genomic island that was likely acquired by horizontal gene transfer.

**Figure 2.**
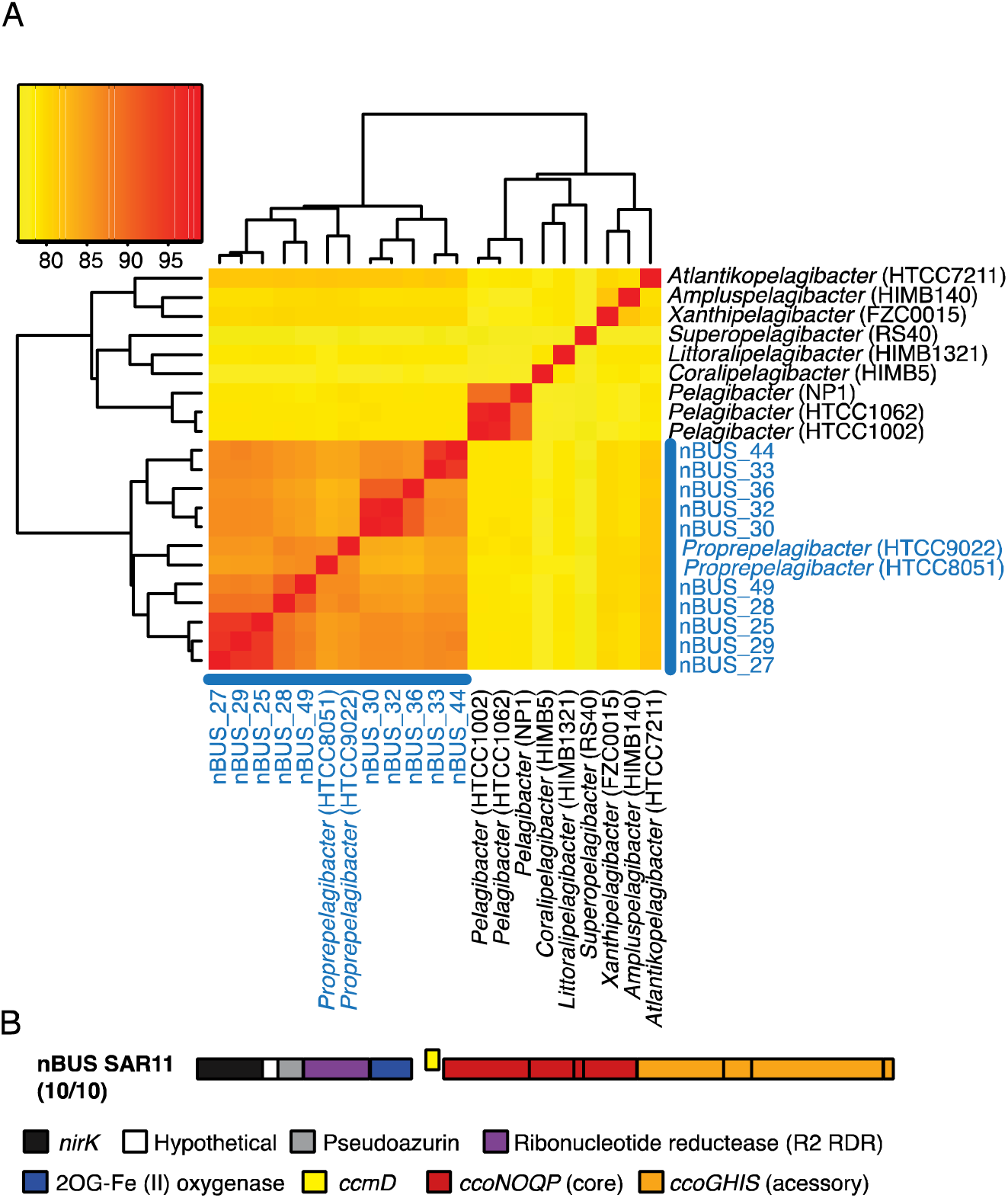
Characteristics of SAR11 isolated from the northern Benguela Upwelling System (nBUS). A) Pairwise average nucleotide identity (ANI) analysis of nBUS SAR11 genomes and the genus *Proprepelagibacter* (blue) and other *Pelagibacter* genera (black), and C) genomic region with denitrification and cbb3-typ cytochrome c oxidase genes in nBUS SAR11 genomes. Labels in panel A correspond to genera designations (Freel et al., 2024) and nBUS isolate IDs. IDs are: *nirK*, copper containing nitrite reductase; *ccm*, cytochrome c biogenesis; and *cco*, high affinity cytochrome c oxidase. Genes correspond to KEGG orthology numbers are listed under gene_id in Tables S2 and S4: *nirK* (K00368, K00362, K00363); *ccoNOPQ* (K00404-K00407).

Genes for the cbb3-type cytochrome c oxidase are arranged in a canonical operon (Decluzeau et al., 2008) that includes the catalytic subunit (*ccoN*), the cytochrome subunits involved in electron transfer (*ccoO* and *ccoP*), and a small subunit with an unknown function (*ccoQ*). This region also has accessory genes *ccoGHIS* (Koch et al., 2000). The four genes between *nirK* and the *ccoN* code for a protein of unknown function, a pseudoazurin, a Fe(II)/2- oxoglutarate oxygenase, and subunit beta of ribonucleotide reductase (R2 RNR). Of these, only pseudoazurins have known roles contributing to denitrification (Fujita et al., 2012).

None of the nBUS SAR11 genomes code for a nitric oxide reductase (NOR). However, Cco proteins share significant homology with NORs, which has led to considerable debate over the evolutionary origin of cbb3-type cytochrome c oxidases, especially regarding their roles in the evolution of aerobic respiration (Ducluzeau etal., 2008; Gribaldo et al., 2009). This also raises the possibility that some extant cbb3-type cytochrome c oxidases could be bifunctional enzymes that reduce oxygen under oxic conditions and NO to N_2_O under anoxic conditions. It has been known for some time that cbb3 heme copper oxidases can reduce NO to N_2_O in vivo (Forte et al., 2002). Interestingly, two of the genes between *nirK* and cbb3-type cytochrome c oxidase are related to proteins that bind NO. The Fe(II)/2-oxoglutarate clavamate synthase from *Streptomyces clavuligerus* (Zhang et al., 2002) and the R2 protein of ribonucleotide reductase from *E. coli* (Haskin et al., 1995). The R2 protein in *Escherichia coli* not only binds NO but also produces N_2_O in vivo. Regardless, this OMZ adaptation may grant SAR11 the genetic potential to grow under both microaerophilic and anoxic conditions, contributing to nitrogen loss in the nBUS, and potentially other OMZs.

### Conclusion

Genomic evidence obtained in this study suggests that denitrification potential in the nBUS is distributed among a diverse community of ubiquitous and abundant lineages of marine bacteria, including SAR11 and UBA868. No chemoautotrophic SUP05 with the genetic potential to respire nitrogen oxides were cultured in this study. However, most of the bacteria isolated and sequenced in this study have the genetic potential to contribute to the initial steps in denitrification, either nitrate or nitrite reduction (**Figure 1**). Notably all organisms with genes to respire nitrate or nitrite also have genes for cbb3-type cytochrome c oxidases, indicating the potential to respire oxygen at low concentrations. Relatively few have the genetic potential to contribute to more than one step in denitrification and only one isolate was identified with the genetic potential for complete denitrification (*Pseudopelagicola* nBUS_20). Evidence that SAR11 have genes to reduce nitrite to NO, and possibly NO to N_2_O under some conditions_,_ suggests that they could be a critical link in denitrification. Similarly, members of a ubiquitous family of sulfur oxidizing mixotrophs in the order *Arenicellales* (Balter et al., 2023) have the potential to respire nitrate and nitrite. This suggests that genome flexibility is a key to the success of these globally distributed marine bacteria.

## Materials and Methods

### High-Throughput Cultivation

We used a high-throughput dilution to extinction approach to isolate bacteria from a low dissolved oxygen (24 µmol/kg) sample collected on April 15, 2024 at a depth of 225 m in the BUS (22°59’58.2"S 13°30’E), as previous described (Monaghan et al., 2020). Briefly, 1 ml of seawater was amended with filter-sterilized glycerol to a final concentration of 10% (v/v), flash frozen in liquid nitrogen, and stored at -80 °C. The sample was later thawed on ice and cells were diluted to 5 and 50 cells/ml in 30 kDa filter-sterilized Puget Sound seawater and amended to 100 µM thiosulfate. Two mL of the diluted cell suspension was then added to each well of a 96 well Teflon plate and incubated at 13 °C in the dark for four weeks. Wells were screened for growth each week using a Guava easyCyte flow cytometer (Cytek, Fremont, CA). Bacteria in wells that were positive for growth (>10^5^ cells/ml) were transferred to 1 L acid-washed and autoclaved polycarbonate bottles containing filter-sterilized seawater amended to 100 µm thiosulfate and monitored for growth until they reached between 10^5^ and 10^6^ cells/ml. Cells were then filtered onto 47 mm 0.2 µm isopore membrane filters (MilliporeSigma, Burlington, MA) and stored at - 80 °C.

### High Molecular Weight DNA Extraction

Filters with cells from cultures were placed in sterile 2 mL DNA LoBind tubes (Eppendorf, Hamburg Germany) and cut into smaller pieces with sterile scissors. Sterile TE buffer (200 µL) was added, and filters were frozen at -80 °C for 20 minutes and then thawed at room temperature. High molecular weight DNA was extracted from the filters using a modified Autogen Quick Gene DNA Tissue Kit (Autogen, Holliston, MA), modified by doubling all recommended extraction volumes and eluting DNA in 200 µL of molecular grade water. DNA was purified using 1X v/v DNA magnetic beads (Sergi Lab Supplies, Seattle, WA) with two 80% ethanol washes then eluted in 20 µL of molecular grade water. Purified DNA was quantified using a Qubit dsDNA Quantitation High Sensitivity kit (Invitrogen, Waltham, MA).

### Long-read Genome Sequencing and Assembly

High molecular weight DNA libraries for whole genome sequencing were prepared using the SQK-RAD114 rapid library prep kit (Oxford Nanopore Technologies, Oxford, UK) and sequenced on a Minion equipped with Flongle R10.4.1 flow cells. DNA bases were called using the Dorado basecalling algorithm (dna_r10.4.1_e8.2_400bps_hac@v4.3.0).

Genomes were assembled from fastq files with Flye v2.9.1 (Kolmogorov et al., 2019) and polished with Medaka 1.7.2 (github.com/nanoporetech/medaka) using the galaxy web platform (Community, 2024). Single circular contigs were evaluated using DFAST for the initial identification and characterization (dfast.ddbj.nig.ac.jp)(Tanizawa et al., 2018).

### Genome Annotation and phylogenomic analyses

The final annotation was completed by NCBI using the Prokaryotic Genome Annotation Pipeline v6.8 (Tatusova et al., 2016; Li et al., 2021). Bacterial taxonomy was determined using GTDB-Tk v2.3.2, the de-novo workflow and the v214 reference database (Chaumeil P-A, et al., 2022). All isolates were given the prefix “nBUS” followed by the cultivation number. SAR11 species were identified by average nucleotide identity (ANI) with Pan-genome Explorer (Dereeper et al., 2022; Perrin and Rocha, 2021). Phylogenetic trees were constructed with MUSCLE v3.8.31 (Edgar RC. 2004) and RAxML v8.2.11 with model GTRGAMMA (Stamatakis A, 2014) using the ETE3 v3.1.3 phylogenetic analysis pipeline (Huerta-Cepas J, et al., 2016). Whole genome phylogenies were constructed using the bacterial_71 single-copy core gene collection in anvi’o v8 (Eren et al., 2021). Genomes were annotated for comparative analysis of denitrification genes using DRAM (ver. 1.1.1) (Shaffer et al., 2020) with the KEGG (Kanahisa et al., 2023), UniRef90 (Suzek et al., 2015), PFAM (Mistry et al., 2021), dbCAN (Yin et al., 2012), and MEROPS (Rawlings et al., 2018) databases.

## Supplementary Material

Supplementary material is available at *Genome Biology and Evolution* online.

## Acknowledgements

We would like to thank the captain and crew of the R/V Marcus G. Langseth and the chief scientist and science party for help with sample collection (MGL24-05).

## Data Availability

Genome sequences were deposited in GenBank under Bioproject PRJNA1173757 with accession and sequence read archive identifiers for each genome (**Table S1, Supplemental Material**).

## Author Notes

Robert M. Morris and Timothy E. Mattes contributed equally to this work.

